# Physiological and functional characterization for high-throughput optogenetic skeletal muscle exercise assays

**DOI:** 10.1101/2025.06.02.657505

**Authors:** Ronald H. Heisser, Angel Bu, Laura Schwendeman, Tamara Rossy, Ritu Raman

## Abstract

Exercise has long been considered an essential part of human health and longevity. Recent physiological studies have expanded muscle’s role beyond simply acting as an actuator, revealing muscle’s exercise-mediated paracrine and endocrine relationships with other organ systems. *In vitro* engineered skeletal muscle models can address physiological questions about exercise adaptation with the precision of cell biology. Optogenetic tools have enabled a noninvasive approach to stimulating muscle contraction that avoids the potential off-target effects of electrical stimulation techniques. In this article we propose high-throughput culture and optical exercise protocols to generate statistically robust cellular exercise response datasets. We characterize optical rheobase for 2D muscle tissue morphology, finding that optical intensities as low as 5 μW mm^-2^ can trigger contraction. We then analyze bulk RNA sequencing data collected from high throughput, acute exercise protocols and find a rich display of transcriptional behavior that is consistent with experimental observations. The spontaneous contractility of our tissue constructs introduced oxygen diffusion challenges when maintained in a 24 well plate, and our analysis shows divergent myogenic and pathological transcriptional consequences of hypoxia. We believe our techniques provide a practical foundation for conducting future high-precision *in vitro* exercise studies of skeletal muscle.

**Translational impact:** High-fidelity, engineered skeletal muscle has potential to elucidate exercise-mediated response pathways at the cell and tissue level. We leverage optogenetic techniques to develop a high-throughput assay that optically stimulates 2D muscle monolayers, avoiding potential cell damage from electrical stimulation. Our culture and exercise protocol generates statistically robust RNA sequencing datasets which reveal myogenic and pathological responses to exercise and *in vitro* culture conditions, informing practical next steps to cultivate stronger, more physiologically relevant muscle models.

## Introduction

Appreciation of exercise’s physiological implications must grow in a technologically modern society that continually reduces our need to move our bodies^1,2^. Mobility is primary to our mammalian experience and a critical indicator of longevity in human health^3^. Exercise can sustain a level of skeletal muscle composition necessary for bodily resilience (such as impact from falls), strength, and cardiovascular health^4^. Beyond a biomechanical role, recent research has implicated muscular movement in biochemical signaling with many organ systems. While glucose uptake and thermogenic pathways have been understood for decades, exercising muscles also release signaling molecules, “myokines”, that can control vascularization, bone formation, immune system function, peripheral nerve growth, and even GLP-1 secretion^5–8^.

*In vivo*, motor neurons electrochemically stimulate muscle contraction at the neuromuscular junction (NMJ), triggering a precise depolarization cascade through the tubular sarcoplasmic reticulum that releases intracellular Ca^+2^ to sarcomeres^9^. The same excitation-contraction coupling has not been efficiently recreated via artificial means, though some NMJ co-culture models have produced measurable muscle contractions^10–15^. In laboratory settings, muscle contraction can be controlled using electrical and optical stimulation techniques, although excitation pathways may differ from native tissue^16,17^.

Tissue engineering offers a suite of biofabrication tools to study *in vitro* skeletal muscle function, enabling cellular visualizations (such as muscle fiber width and contractile displacement) and characterizations that illustrate less-visible physiological processes (like myokine release) in isolation^18,19^. Therefore, it is important to develop tissue constructs with a similar contractility to native skeletal muscle. Using fibrin, collagen, and Matrigel-based hydrogel ECM-mimicking substrates, tissue engineers can currently produce muscle constructs with up to 10% of the contractility of *in vivo* muscle^20^.

Research employing responsive, contractile engineered muscle constructs has successfully emulated the hypertrophic and myogenic effects of exercise^21,22^. Most studies use electrical stimulation because it can be simply implemented by placing electrodes in media and controlling voltage inputs via a function generator.^23,24^ However, electrical stimulation risks electrolyzing media, causing fiber detachment, and electroporation at improper input parameters^19,25^. Additionally, electrical stimulation can depolarize voltage-gated channels not associated with contraction, potentially inducing off-target transcriptional responses.

By contrast, optogenetic stimulation non-invasively targets channelrhodopsin (ChR) membrane proteins. Light can be precisely timed, like electricity, but also has the advantage of spatial patterning that enables it to differentially stimulate smaller groups of muscle fibers within a single tissue construct, more closely resembling motor unit control. We have previously shown >5x increase in force production in optogenetic muscle using 1 Hz 30 min optical exercise regimens^26,27^. Furthermore, building on work from others showing optogenetic stimulation enables non-invasive control of muscle contraction *in vivo*^28^, we have demonstrated that stimulation of optogenetic muscle grafts implanted *in vivo* enables functional recovery from volumetric muscle loss injury in mice.^29^

Optical stimulation has clear advantages when considering constructs that contain more than one muscle in a volume of media. Multi-muscle models can more closely model *in vivo* conditions, like antagonistic behavior, but are hard to electrically isolate, making electrical control implementation more difficult. *In vivo* studies note that stimulating the correct muscle, when surrounded by several other muscles and organs, is easier with optical stimulation^30^. Optical stimulation also has the potential to simplify high throughput experiments, given that well plates usually have transparent bottom surfaces and lights can easily be patterned to match 24, 96, and 384 well plate form factors.

Given the potential advantages of optical stimulation *in vitro*, there is a need to understand and characterize the relationship between repeated light exposure, muscle contraction responses, and longer-term metabolic responses. Studies have previously characterized light stimulation parameters for muscles from ChR2-expressing mice using explanted tissues and transcutaneous optical stimulation^28,31^. The variable light absorption of 3D tissues and skin complicates a fiber-level understanding of optical contractility. *In vitro*, 2D optogenetic studies quantify more fundamental qualities such as depolarization currents and maturation indicators (fusion index, myotube diameter, etc.), but use constructs with contractions (<10 μm displacements) that may not be strong enough to induce the full breadth of exercise responses in muscle cells, even at mm tissue scales ^32–34^. Another challenge to 2D monolayer fidelity is the tendency of fibers to delaminate from their substrate during differentiation, due to incompatible substrate stiffnesses, poor cell adhesion features, and contractile force development^35–37^.

Utilizing previously developed tissue protocols^7,38^, we cultured highly contractile 2D muscle constructs (with mean average displacements up to ∼100 μm) on micropatterned fibrin gels and characterize their twitch contractile response by varying optical intensity, frequency, and pulse width. We established minimum intensity thresholds (rheobase) for inducing twitch, quantities that, to our knowledge, are not currently documented in optogenetic *in vitro* muscle literature. Recent literature has identified contractile dysfunction stemming from ChR2-EYFP overexpression in mice, leaving additional questions of potential downstream effects in optogenetic engineering strategies^39^. We use ChR2-expressing C2C12 mouse myoblasts as the model system for this study because it is among the most widely used cell lines in skeletal muscle engineering, producing meaningful physiological tissue models as well as biological actuators for robotics^40,41^. We focus on the ChR2(H134R) protein variant because it has proven to efficiently activate muscle contractions in multiple tissue engineering contexts, including biohybrid robotics, but future studies may need to perform optimization on other optogenetic ion channels to determine cross-ChR relevance^26,28,42^. As optogenetic exercise training involves repeated light exposure over days or weeks of culture, we conduct high throughput optical exercise experiments and bulk RNA sequencing (RNA-seq) transcriptomic analysis to examine optical dose influences on metabolic and contractile function. We find robust contractile, metabolic, and extracellular transcriptional responses to exercise that characterize deviations from native physiological states, motivating developments to better model exercise in highly contractile tissue engineered muscle constructs and to tune development protocols to manufacture stronger biohybrid robots.

## Results

### Optical excitation-contraction characterization

We recorded and analyzed contractile responses from light pulses of varying intensity to determine the effective bounds of functional optical dose, setting context to evaluate downstream effects of repeated stimulation in our exercise protocols. C2C12 myoblasts expressing ChR2(H134R) light-sensitive proteins grew into differentiated 2D muscle sheets on micropatterned fibrin gels, evenly exposing individual cells to light stimuli. Fig 1a shows the 6-well plate format used in this study. The cell-fibrin construct sits in a 15 mm-diameter acrylic well (similar to that of a 24-well plate) to limit tissue size. This design also allowed us to have ample culture medium with a minimal depth of liquid above the cells, facilitating better oxygen exchange with surrounding air^43^. To align myoblasts during differentiation, we leveraged our established protocols to create 25 μm parallel grooves in the fibrin gels as they cure^38^. Figs. 1b and 1c respectively show brightfield and fluorescent images of our 2D tissues.

**Figure 1:**
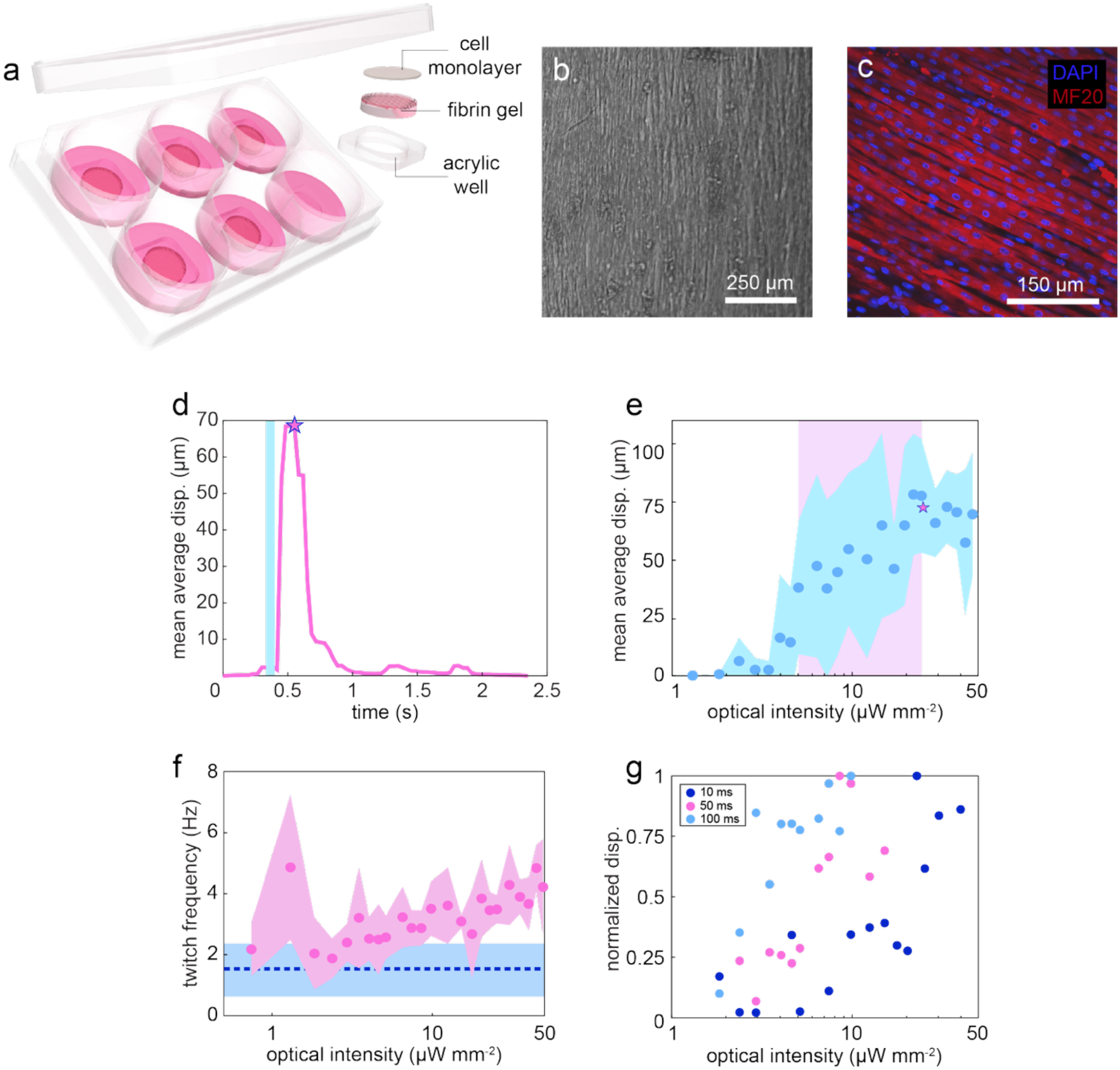
Optical excitation-contraction coupling characterization. a) 6 well plate design. Cells are seeded on fibrin gels in acrylic wells. b) Brightfield microscope image of aligned muscle fibers at 4x magnification. c) Myosin (MF20) and nuclear (DAPI) antibody stains of muscle fibers. d) Contractile response from a light pulse with an intensity of 25 μW mm^-2^. The blue band shows the duration of the optical pulse. e) Average values of optical response curves. The pink shaded region shows the range of rheobase measured across samples. f) Average twitch frequency of samples under continuous illumination. Dotted line and shade correspond to average and standard deviation of recorded spontaneous twitch. g) Normalized optical response of a single tissue sample exposed to varying optical pulse widths, showing a trend of lower rheobase for longer optical pulses.

We obtained optical excitation-contraction response curves over all biological replicates by testing 50 ms light pulse durations and intensities between 0.75-50 μW mm^-2^. 50 ms pulse widths have been used in previous optogenetic exercise studies and help to distinguish stimulated contractions from spontaneous contractions at lower intensity ranges^33^. We used a collimated LED to spot-illuminate tissues, recording videos at 30 fps using a microscope camera (Fig. S1), finding peak displacement amplitudes for each video (Fig. 1d, Movie S1). We observed that optically stimulated contractions became perceptibly distinct from spontaneous contractions at intensities as low as 3 μW mm^-2^, and that all tissues responded at thresholds below 20 μW mm^-2^. This power range is lower than other 2D optogenetic muscle experiments in literature, reporting light intensities similar to those used during in vivo experiments (0.1-1 mW mm^-2^)^28,32,34,44^. Fig 1e shows an increasing contractile response that, on average, plateaus around 25 μW mm^-2^. Because tissues display expected variability in optical sensitivity between replicates and individual muscle fibers, precision characterization likely involves calibration stimulation parameters to each tissue and experimental setup. Our initial experiments suggest that rheobases in 2D C2C12 tissues, at 50 ms pulse durations, are in the range of 5-25 μW mm^-2^.

We note that at our relatively low light intensities, our 2D *in vitro* tissue constructs respond to continuous optical exposure differently than native tissues; Bruegmann et al. report that continuous illumination causes muscle to briefly contract, eventually returning to a relaxed state. Continuous stimulation in our tissues mimicked the dynamics of spontaneous contraction but with an increased frequency of twitching (Fig. 1e, Movie S2). At higher light intensities we observed that tissues would briefly switch to tetanic contractions, therefore we anticipate that beyond our stimulation parameters, more tetanic behavior would be observed^45^. An uptick in twitch dynamics was apparent in continuous stimulation at low intensities (2 μW mm^-2^), showing some baseline optical contribution to otherwise cell-generated membrane depolarizations. We also observed this behavior when examining and filming tissue contractility with white light brightfield illumination, where the difference in twitch frequency and magnitude was more pronounced. Illuminating tissues with red light avoids this background experimental influence. We used red light illumination for our optical characterization study. Fig. 1f compares contractile responses for variable duty cycles, showing the tradeoff between threshold stimulation intensity and duty cycle. Selecting the optimal twitch stimulation parameters likely involves balancing optical dose (the product of intensity and duty cycle, given in nJ) and aggregate channel opening dynamics. From Fig 1f, our minimum optical dose ranged 200-300 nJ, in agreement with reported electrical pulse energy values from an earlier rheobase characterization study^25^.

Twitch characterization helps determine optimal short-term stimulation parameters (intensity, frequency, duty cycle) for triggering contraction. Intuitively, minimizing the optical dose minimizes potential cellular phototoxicity, yet tissue contractility tracked over the course of an experiment may not fully reflect a cell’s metabolic state. We considered the longitudinal effects of optical stimulation using bulk RNA-seq transcriptomic analysis in the next section.

### High-throughput exercise and transcriptomic analysis

Bulk RNA-seq has potential to give a rich description of systemic cellular activity of exercise conditions in our 2D optogenetic muscle constructs. Investigating the transcriptomic behavior of cells undergoing long-term optogenetic stimulation can also help clarify the effects of extended light exposure on metabolic and regulatory processes, and how muscle-specific physiological adaptations, like fiber-type transformations, can couple influences from exercise and optical stimulation. We cultured muscle constructs in a 24-well plate format and conducted high-throughput, acute exercise protocols by modifying an computer-controlled optical stimulation platform from Repina et al. to match our well plate geometry (Fig. 2a)^46^. This system allowed us to keep cells inside the incubator during exercise and minimize environmental effects.

**Figure 2:**
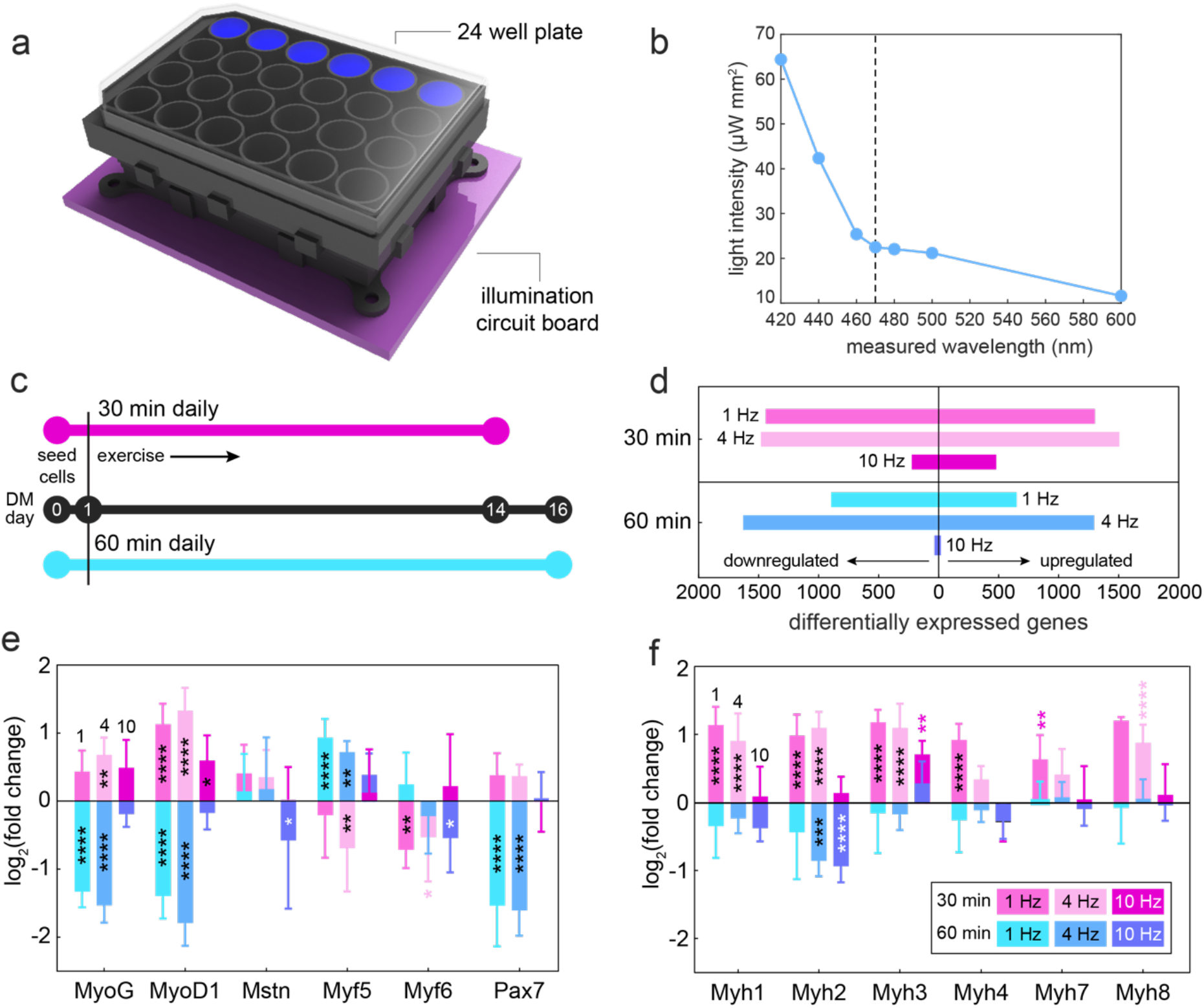
Physiological characterization of optical exercise. a) High throughput optical stimulation platform, adapted from prior work^46^. b) Spectral characterization of stimulation platform. Dotted line shows intensity at ChR2 principal stimulation wavelength we report in this study. c) Tissue culture and experimental timeline for 30 min and 60 min experiments. DM stands for differentiation medium. d) Bar graph of significantly upregulated gene counts (log_2_(fold change) > 0.5, p_adj_ < 0.05) for each experimental condition. e) Measured log_2_(fold change) values for genes associated with muscle differentiation and maturation. f) Measured log_2_(fold change) values for myosin heavy chain (Myh) protein-encoding genes used to identify muscle fiber types. See SI XX for a table of Myh proteins and their associated fiber types.

Conditions were tested at the upper end of our rheobase characterization results, with a light intensity of 22.5 μW mm^-2^, at 470 nm, shown in Fig. 2b. We optically trained sets of six biological replicates alongside an unexercised control condition at 1 Hz, 4 Hz, and 10 Hz stimulation frequencies for 30 min or 60 min per day. These stimulation parameters were previously shown to enhance contractility in 3D optogenetic muscle actuators^26,27^. 1 Hz and 4 Hz stimulation represent low- and medium-exertion conditions, respectively, and the 10 Hz frequency emulates a high-exertion condition that approaches tetanus but remains within the operating limits of our stimulation platform. Cells remained in 600 μL of media throughout experiments and underwent optical stimulation within one hour of media replenishment. Fig. 2c shows experimental timelines for the 30 min and 60 min exercise experiments.

Bulk RNA-seq transcriptome analysis allows us to explore and compare the physiological tissue states from each exercise condition. The 1 Hz and 4 Hz conditions had comparable upregulated and downregulated (log_2_(FC) > 0.5 and p_adj_ < 0.05) gene counts. In both experiments, the 10 Hz condition produced the lowest number of significantly upregulated or downregulated genes (Fig. 2d). In fact, we found that the most significantly downregulated gene in the 10 Hz, 60 min condition is Myh2, the myosin heavy chain protein that identifies type 2a (fast-oxidative) muscle fibers.

Reviewing protein-encoding genes associated with fiber type specification and fiber maturation revealed that the 60 min experiment transcriptionally mirrors the 30 min experiment. 1 Hz and 4 Hz conditions in the 30 min experiment showed significant upregulation of most myosin isoforms, when, coupled with upregulated MyoG, MyoD, and downregulated Myf5, could indicate cellular differentiation processes^47,48^. Given the presence of Myh3 and Myh8, embryonic and neonatal myosin isoforms, many fibers likely remained in an immature state at the time of extraction. Given the presence of Pax7, it is possible that new fibers are also still forming at rates comparable to those undergoing fiber type commitment. While the 60 min experiment displayed significant downregulation of myogenic regulatory factors, the relative parity of Myh transcription (except Myh2) to control matches our observations that 60 min experimental conditions remained contractile and responsive even after a fatigue test.

Interpretative complications can arise when tying cell function to differential expression of individual genes, especially for regulatory genes implicated in multiple pathways. Gene set enrichment analysis (GSEA) abstracts away from transcriptional granularity and maps aggregate differential expression to all dimensions of chemical and phenotypic cell behavior, from mitochondrial complex synthesis to cellular chemotaxis. Fig. 3 explores systemic effects of our exercise protocols between the two 1 Hz experimental conditions using Hallmark and Gene Ontology (GO) gene set collections. The Hallmark collection comprises 50 well-defined gene sets and enables clear cross-experimental physiologic comparisons (Figs. 3a, 3b), while GO annotations capture the hierarchical organization of thousands of cellular processes and help us capture a broad range of cellular activity (Figs. 3c-3g).

**Figure 3:**
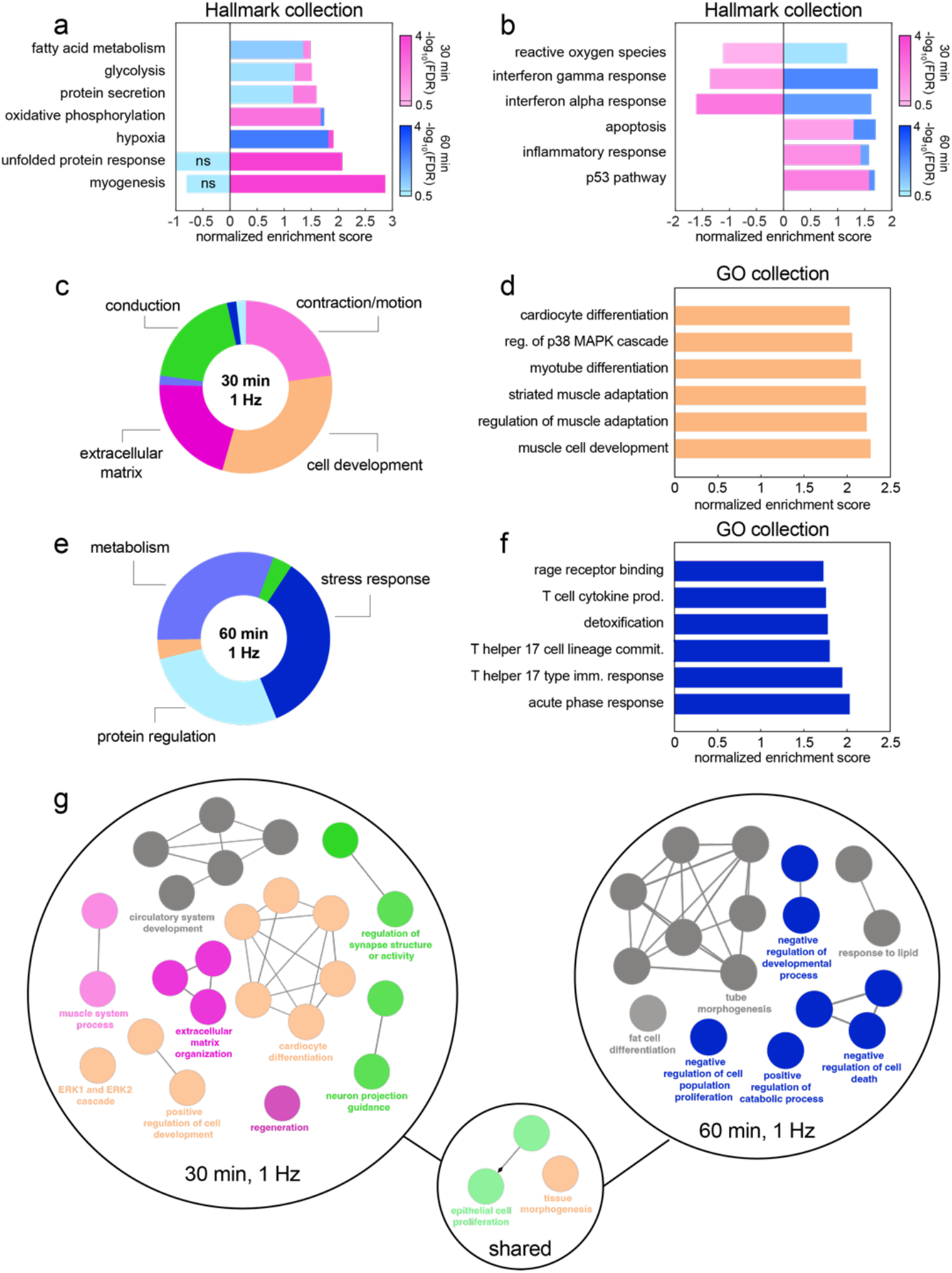
Gene set enrichment analysis (GSEA) results. Cell transcriptional behavior for 1 Hz conditions from 30 min and 60 min experiments are compared using Hallmark and Gene Ontology (GO) gene set collections. Hallmark normalized enrichment scores (NESs) comparing gene sets associated with a) muscle development and metabolism, and b) inflammatory responses in both 1 Hz conditions. Lines across the color bars indicate the cutoff for statistical significance, shown on individual bar plots with “ns”. c) From the 75 most enriched GO terms in the 30 min condition, 59 each fit into one of the seven categories shown in c) and e), centered around conduction, contraction, ECM, and cell development processes. d) Examples of enriched cell development gene sets in the 30 min condition. e) 55 of the 75 most enriched 60 min GO terms mostly represented metabolic, stress response, and nonspecific protein regulation processes. f) Examples of terms representing stress response behaviors in the 60 min condition. g) CytoGO visualization of genesets associated with the 250 most upregulated genes from each condition. Grouped terms are color coded as in c) and e). Gene sets not associated with a category are colored gray. Shared gene sets are centered between the two experimental groupings.

Hallmark genesets show snapshots of functional muscle behavior in Fig. 3a. Myogenesis was the highest-ranked (most enriched) gene set in the 1 Hz, 30 min condition and the lowest-ranked gene set in the 1 Hz, 60 min condition (not statistically significant), aligning with trends observed in Fig. 2. The closely matching upregulation of hypoxia between both conditions matches experimental observations of accelerated media yellowing (indicating reduction of pH) as monolayers became more contractile. Contracting muscle has oxidative metabolism 10-100 times higher than resting muscle^49^, so it is reasonable that even a monolayer of cells can outpaces oxygen diffusion through a column of liquid several mm in thickness. Given the high contractility and occurrence of spontaneous twitch in our monolayers, baseline oxygen consumption is likely higher than expected for resting muscle tissue. Yet, Fig. 3b reveals differences in tissue response to hypoxic and exercise-based stressors. The 30 min condition had significantly downregulated reactive oxygen species and interferon responses (implicated in muscle fatigue and repair), while the 60 min condition was significantly upregulated in all pathways. In general, inflammatory response upregulation can be attributed to multiple causes and play a complex role in muscle cell development and homeostasis^50^. However, the p53 pathway is primarily implicated in managing oxidative stress and has been shown to regulate mitochondrial synthesis in muscle^51,52^.

To explore a wider range of cellular processes, we investigated the 75 highest enriched GO annotation gene sets for each experimental condition and categorized relevant annotation terms into one of the seven categories seen in Figs. 3c and 3e. Other than contraction-related annotations, the 30 min condition displayed significant signals of conduction regulation, extracellular matrix (ECM) remodeling, and cell development behaviors. Conduction-related genesets primarily indicate protein expression related to transmembrane ion transport but also include neuron morphology gene sets, such as dendrite morphogenesis (GO: 0048813). ECM genesets indicate general reorganization and collagen synthesis processes, including one basement membrane organization term (GO: 0071711) that is relevant to muscle. Interestingly, we found that during RNA extraction, the 30 min, 1 Hz condition was the most difficult to scrape away from the fibrin gel, experimentally indicating stronger cell adhesion and ECM activity. Fig. 3d shows examples of enriched cell development genesets, mostly pertaining to muscle differentiation and adaptation. Multiple cardiac contraction, morphogenesis, and development terms are present in GSEA results, yet this can be expected given the set of shared contractile and conductive proteins between muscle and cardiac tissues^53^.

The 60 min, 1 Hz GO annotations presented a large proportion of enriched stress-response gene sets (Fig. 3f) accompanied by significant amounts of metabolic modification and nonspecific protein synthesis. Hallmark gene set analysis indicated that both conditions are under hypoxic stress, yet stress response terms seen in the 60 min condition are not developmental or myogenic like that of the 30 min condition; Fig. 3f shows several annotations more indicative of pathological tissue states. Metabolic gene sets include several subprocesses associated with mitochondria formation and oxidative metabolism.

In addition to exploring individual enriched Hallmark and GO gene sets, we visualized gene set groupings using the ClueGO, a Cytoscape plugin, to capture nonredundant GO groupings (Fig. 3g)^54^. ClueGO visualization indicates similar behavioral trends we classify with our other analyses but presents additional groupings not immediately evident when looking through individual gene sets. One example of this is the presence of several gensets pertaining to renal development, and epithelium formation in both exercise conditions. We did not initially find commonality between these annotations. In the 60 min condition, ClueGO identified hierarchical commonality in tube morphogenesis (Fig. 3g, 60 min condition, GO: 0035239), which comprises general processes associated with tube formation across organ systems, including renal, digestive, and circulatory system epithelia. ClueGO also identified 30 min condition had a large set of interrelated gene sets corresponding to circulatory system development. We believe that in both cases, cells released cytokines or made efforts to adapt themselves to address their hypoxic environment. While further investigation is needed to determine with detail the specific cellular behaviors, we note that the Hallmark angiogenesis gene set is significantly enriched in the 30 min condition, matching the circulatory system development (GO: 0072359) grouping in Fig. 3g.

## Discussion

Optogenetic culture and stimulation techniques offer a non-invasive method of stimulating excitation-contraction couplings in *in vitro* engineered muscle tissues. When coupled with high-throughput culture formats, the combination offers a simple, powerful means to gain statistically robust functional and physiological data from long term exercise experiments. Further, protein-specific optical stimulation reduces potential for off-target cellular responses in comparison to electrical stimulation. We previously developed substrate biofabrication strategies for aligned 2D muscle constructs in well plate form factors but did not quantify relationships between stimulation parameters and contractile magnitude. This is especially important for optical stimulation given the binary optogenetic contractile response in 2D morphologies. Our characterization experiments show that rheobase is achieved at light intensities between 5-25 μW mm^-2^, one to two orders of magnitude lower than intensities used in other 2D optogenetic studies.

One consequence of growing highly contractile muscle tissue *in vitro* is a consistent behavior of muscle cells to routinely depolarize their own sarcolemma and initiate contractions, perhaps counterintuitively, in the absence of motor neuron innervation. Spontaneous contraction is now understood as a necessary step in myogenesis, where spontaneous contractions complement passive tension development, aligning contractile and structural proteins during sarcomere synthesis^55^. Further, spontaneous contractions provide the necessary signals for embryonic NMJ formation^56,57^. Even adult muscle, when denervated, has potential to develop spontaneous contractions^58^. Numerous 2D and 3D *in vitro* studies have described this phenomenon with varying degrees of interest^37,59–61^.

While we mention spontaneous twitch throughout this article, we believe it is a significant consideration for our 2D optogenetic morphologies, given their displacement magnitudes at this scale. Spontaneous contractions complicate the functional and physiological characterization of tissue exercise protocols. For example, the self-generated depolarization patterns are coordinated between groups of fibers and can interfere with the repeatability of optical contraction initiation. During occasional bouts of vigorous twitching, cells seemed resistant to optical pulse interruptions of their self-established contraction rhythms, requiring a waiting period before cells became receptive to optical control. Additionally, the consistency of regular twitching we observe upon removing cells from the incubator (even when turning ambient lights off) indicates that these cells likely twitch throughout day and night. This implies that cells effectively exercise themselves and can induce their own physiological responses independent of a given exercise protocol. We assume that “control” conditions are unexercised, but spontaneous contractions may decrease differential expression and contractility between control and exercise conditions, especially for acute exercise protocols.

Since ChR2(H134R) is a voltage-gated membrane channel protein, it may modulate spontaneous twitch characteristics, interesting to study in later investigations. Instead of delivering light pulses to stimulate contraction, future exercise protocols can embrace spontaneous twitch behavior by delivering low-intensity, sustained amounts of light, modulating twitch frequency (see Fig. 1g). We note that spontaneous twitch does decrease over time when well plates are left outside the incubator and media cools to room temperature and as media becomes used, but these methods cannot be deployed to study tissue function in physiologically relevant circumstances.

Despite the presence of spontaneous twitch, our optical exercise protocols still produce meaningful differential gene expression among the 6 experimental conditions we test. RNA-seq transcriptomic data reveals similar trends in cell behavior when comparing individual genes (Figs. 2e, 2f) and GSEA results (Fig. 3). Overall, transcriptomic analysis points to myogenic responses in the 30 min experiment (particularly the 1 Hz and 4 Hz conditions) and pathological stress responses in the 60 min experiment. Discussing “stress response” in exercise contexts is difficult because exercise itself is a form of stress. Hypoxia is another form of stress; our observations of accelerated media yellowing rates and reduced contractility in quiescent (oxygen diffusion gradients undisturbed by agitation) day-old media in all conditions is validated with the enriched Hallmark hypoxia gene set. Yet, like exercise, hypoxia has a multidimensional role in exercise contexts, being itself a consequence of exercise, thus “expected” by muscle to occur periodically. At the right magnitude and duration, hypoxia supports hypertrophy, fiber type transitions, and myogenesis. The tube morphogenesis and circulatory gene sets enriched in both conditions make sense in a hypoxic context, with cells releasing angiogenic cytokines and making attempts to locally adapt to extended hypoxic conditions.

Other signaling pathways enriched in our analysis have indeterminate physiological roles. The Hallmark unfolded protein response (UPR) gene set refers to endoplasmic reticulum activation (here, in the sarcoplasmic reticulum) in response to the accumulation of unfolded proteins, usually a result of cellular stress. We find that the UPR gene set is also highly enriched in the 30 min condition. When coupled with myogenesis, this could indicate myogenic adaptations to stress from exercise or increased fiber commitment to differentiation^62^. Processes downstream from UPR-upregulation can involve selective apoptosis of myoblasts deemed incompatible for differentiation, which could explain signal towards upregulation of the 30 min apoptosis gene set, though not statistically significant. Interestingly, UPR is (insignificantly) downregulated in the 60 min condition despite the apparent state of stress.

Given the complex role of signaling pathways in managing exercise-induced muscle growth and homeostasis^63^, tracking other experimental quantities such as oxygen or glucose consumption over time can illustrate long term adaptations to experimental conditions. Further, the autocrine effects of myokines can accumulate in static media and cause unintended deviations from *in vivo* extracellular chemical conditions. Chronic stimulation exercise protocols generally produce more substantial muscle maturation and will require solutions for maintaining permissible oxygen concentrations^64,65^. Therefore, additional equipment is likely needed to access these longer exercise durations stirring or media automation which can be cumbersome in 24 and 96 well formats^37,66^. Fortunately, optical stimulation platforms illuminate cells outside of and under well plates, leaving room above for control elements to be integrated, unlike electrical stimulation platforms.

Fortunately, the light intensities we characterize and use in this experiment do not seem to directly cause phototoxic cell responses across all experimental conditions. The Hallmark UV response up gene set represents genes upregulated in response to UV exposure. We find that UV response up is enriched in both 10 Hz gene sets (rank 9/50 in the 30 min condition and 10/50 in the 60 min condition), understandable given the relatively extreme stimulation condition. Electrical stimulation studies that use 10+ Hz frequencies do so with bursts of 10 Hz stimulation followed by a brief rest period lasting between 0.4 s and 5 s^23,67,68^. Critical optical dose rates may lie below what we employed in the 10 Hz conditions. Therefore, we believe our stimulation parameters are suitable for chronic stimulation studies, important for inducing longer term maturation, ECM remodeling, hypertrophy, and fiber type shift exercise responses^23,66,69,70^.

## Conclusions

In this article we have built upon recently established 2D engineered skeletal muscle techniques to investigate their potential to faithfully recapitulate exercise-mediated physiological responses in a high-throughput experimental context. We chose to use optogenetically engineered C2C12 myoblasts because of their experimental handling advantages and potential to avoid the pitfalls of electrical stimulation for longer culture timelines. We characterized elemental optical responses, finding that light intensities under 100 μW mm^-2^ are sufficient to stimulate a contractile response. Then performing exercise protocols under these stimulation parameters, we explored physiological cell adaptations using RNA-seq transcriptomic analysis. The divergent transcriptomic profiles of the 30 min and 60 min exercise conditions capture a complex interplay between hypoxia, spontaneous contractility, and the exercise durations of our tissues. Enriched stress response gene sets illustrate pathways that can be interpreted as myogenic or pathological because of the multidimensional roles many of these signaling pathways play during each stage of muscle maturation. The optogenetic characterization and high-throughput tissue culture techniques we present here provide a practical foundation for future high-precision physiological assessments of *in vitro* 2D muscle tissues. Lessons gained from this work inform not only research directions for clinical applications but also the development of stronger biohybrid actuators, where engineered skeletal muscles will be driven at their physiological limits to power robots and other biodegradable machines.

## Materials and methods

### Culture substrate preparation

Culture substrates were prepared similar to prior work^38^. For the optical characterization experiment, 6 well plates were fitted with 15 mm-diameter acrylic wells. Fibrin hydrogel prepolymer was prepared by first mixing a solution of DMEM (Sigma Aldrich, D6429) with 8 mg/mL fibrinogen from fetal bovine plasma (Sigma Aldrich, F8630-1G). Then, 4 μL/mL thrombin from bovine plasma (Sigma-Aldrich, T4648-1KU) was added to the solution, mixed, and pipetted into the acrylic wells at 850 μL per well. As the fibrin gel cured, stamps with 25 μm groove widths were aligned with the acrylic well and placed at the top of the gels to replicate the groove pattern on the gel surface.

Stamps were prepared using prior fabrication methods previously described^38^. Prior to gel insertion, stamps were sterilized overnight in 70% ethanol and with UV radiation the day of fibrin preparation. To prevent adhesion to fibrin, the stamps were first soaked for 15 min in bovine serum albumin (BSA, Thermo Fisher Scientific, 37525) diluted to a 5% vol/vol concentration. Immediately before gel insertion, stamps were removed from BSA and lightly dried with an aspirator to minimize the BSA layer. Then, a small amount of phosphate-buffered saline (PBS, Thermo Fisher Scientific, 20012027) was placed on the stamp surface to coat the grooves and minimize trapped air when submerged in the fibrin prepolymer. Gels cured for 1 hr at room temperature and were maintained in DMEM after stamp removal. Stamps were then cleaned with soap and water using a thin-bristle brush to de-soil the stamp face and sides, then undergoing sonication in 70% ethanol for 30 min. For the exercise experiments, 24 well plates were prepared identically to the 6 well plates except that no acrylic well was used and each well had 500 μL of fibrin.

### Cell culture

Optogenetically engineered C2C12 mouse myoblasts (expressing 470 nm blue-light sensitive Channelrhodopsin [ChR2(H134R)] tagged with tdTomato) were used for all experiments^26^. Cells were expanded in growth medium (GM) consisting of fetal bovine serum (1 : 10 vol/vol, Life Technologies, A5670701), Corning™ L-glutamine solution (1:100 vol/vol, Fisher Scientific Co LLC, MT25005CI), and 1% penicillin–streptomycin (Fisher Scientific Co LLC, MT30002CI) dissolved in DMEM. The cells were seeded on stamped gels at a density of 100,000 cells per well (for both 6 and 24 well plates) and grown in growth medium (GM) with 6-aminocaproic acid (ACA) (Sigma-Aldrich, A2504-100G) at a ratio of 1:50 vol/vol for one day until they reached confluency. They were then switched to differentiation medium (DM) consisting of horse serum (1 : 10 vol/vol, Gibco, 26050-088), Corning™ L-glutamine solution (1:100 vol/vol, Fisher Scientific Co LLC, MT25005CI), and 1% penicillin–streptomycin (Fisher Scientific Co LLC, MT30002CI) in DMEM (Sigma Aldrich, D6429), supplemented with insulin-like growth factor1 (IGF-1) (1:20,000 vol/vol, PeproTech, 100-11R3-1MG) and ACA. Cells in 6 well plates were maintained in 4 mL of media and cells in 24 well plates were maintained in 1.5 mL of media for both GM and DM culture.

### Optical characterization experiment

Cells were seeded in a 6 well plate and cultured for 9 days in DM without exercise before undergoing characterization. On DM day 9, optically stimulated cell contractions were filmed using an Axiocam 202 mono camera (Zeiss) connected to an inverted brightfield microscope (Primovert, Zeiss). To produce the optical pulses, a function generator (Thorlabs) modulated input signals from an LED driver (Thorlabs) to a collimated fiber-optic LED system, previously described^71^. The collimating lens (Thorlabs) was inserted sideways into a fused deposition modeling 3D printed fixture that reflected light downwards to the tissue to avoid collisions with the microscope’s overhead lamp and maintain spotlight perpendicularity. The fixture included alignment elements which centered the light over each well. Optical intensity was measured with a photometer (XXXX Thorlabs) calibrated to the same distance from the light as tissues in the well plate. Plates were illuminated with a 350 lumens red LED hunting flashlight (X.YShine) to avoid unintended stimulation. To gather data, videos were taken at 30 fps of contractile responses to single light pulses while sequentially increasing the intensity, repeating for each well. Displacement fields and mean average displacement approximations of tissue contraction were generated from video frames using an open source tracking algorithm, described previously^72,73^. A custom MATLAB script, specifically utilizing the “findpeaks” function, to determine maximum displacement amplitudes for twitch and stimulated contractions used in Fig. 1. One caveat of this analysis is that the optical pulses briefly saturate the camera, rendering the tracking software unable to track illuminated frames. Since we are interested in what happens after illumination, we track frames backwards in time up to the last frame of illumination to ensure tracking continuity.

### Exercise study

All conditions were exercised daily, starting DM Day 1. Twos open-source optical illumination plates were constructed and kept inside an incubator to facilitate high throughput stimulation^46,74^. The light guide attachments were adjusted in CAD (SolidWorks 2021) to fit the dimensions of our well plate (Cellvis: #P24-1.5H-N). Two 24 well plates were used for each of the 30 min and 60 min experiments, each well plate having two rows filled with a single condition. The illumination platform saves stimulation settings on an integrated Raspberry Pi Zero W computer, so settings were loaded once through the included “JarJarBlinks” Java applet GUI. Media was changed daily, no more than 15 min before exercise.

### RNA sequencing

RNA extraction was performed on day 14 of muscle differentiation for the 30 min experiment and day 16 for the 60 min experiment using the Qiagen RNeasy Mini Kit (Hilden, Germany) following the manufacturer’s instructions. Cell monolayers were treated with the lysis buffer for 15 minutes before being spun down in QIAshredder spin columns to homogenize the tissue.

Sequencing libraries were generated using the NEBNext® Ultra™ II Directional RNA preparation with poly(A) selection kit. Library sizes were quantified and verified via qPCR and Fragment Analyzer before being loaded on the Singular G4 platform in a 50-base paired-end configuration with 8+8 nucleotide indexes.

Reads were mapped against mm39 (GRCMm39) using the BWA-MEM algorithm^75^. Quality control metrics such as mapping rates, unique 20-mers and fraction of ribosomal RNAs were calculated using bedtools version 2.30.0. FASTQ files were processed using the nf-core/rnaseq version 3.11.1 pipeline. We utilized the GRCm39 reference genome and ENSEMBL GRCm39 murine annotations.

Differential gene expression was done in DESeq2 in R version 4.2.0 using the apeglm method to provide Bayesian shrinkage estimators for effect sizes. We reported log_2_ fold changes and Benjamini-Hochberg adjusted p-values. Our Gene Set Enrichment Analysis (GSEA) was done in version 3.0. We utilized the log_2_ fold changes and p-values from DESeq2 to rank the genes in preparation for analysis in the GSEAPreranked module. Two major MSigDB collections were used to investigate transcriptional differences, Hallmark and C5 (GO) gene set collections. We reported normalized enrichment scores from the GSEA analysis with the associated FDR q-values. Gene sets with FDR q-values < 0.25 were considered statistically significant.

## Acknowledgements

The authors acknowledge funding support from the DoD Office of Naval Research Young Investigator Program (N00014-24-1-2060), the DoD Army Research Office Early Career Program and PECASE (W911NF-22-1-0126), and an NSF CAREER Program award (2238715). We thank Vincent Butty for assistance with RNA-seq and Jeffrey Kuhn for helpful discussions.

