## Supplemental Information for "Physiological and functional characterization for high-throughput optogenetic skeletal muscle exercise assays"

**Supplementary Information:**

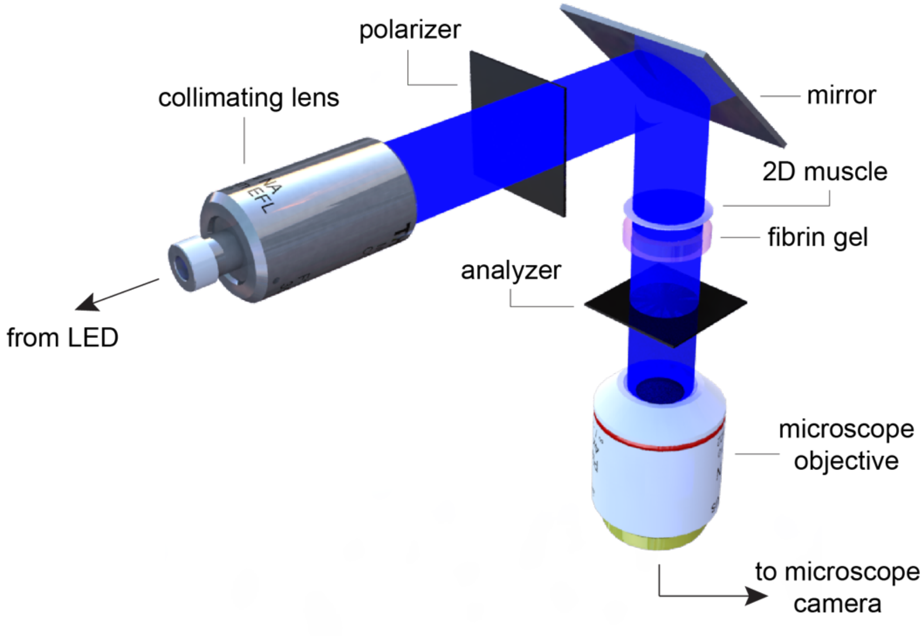


**Figure S1:** Optical characterization setup. Light was polarized to reduce the relative illumination to the camera. Not shown: red LED light source, well plate, and mirror holder.

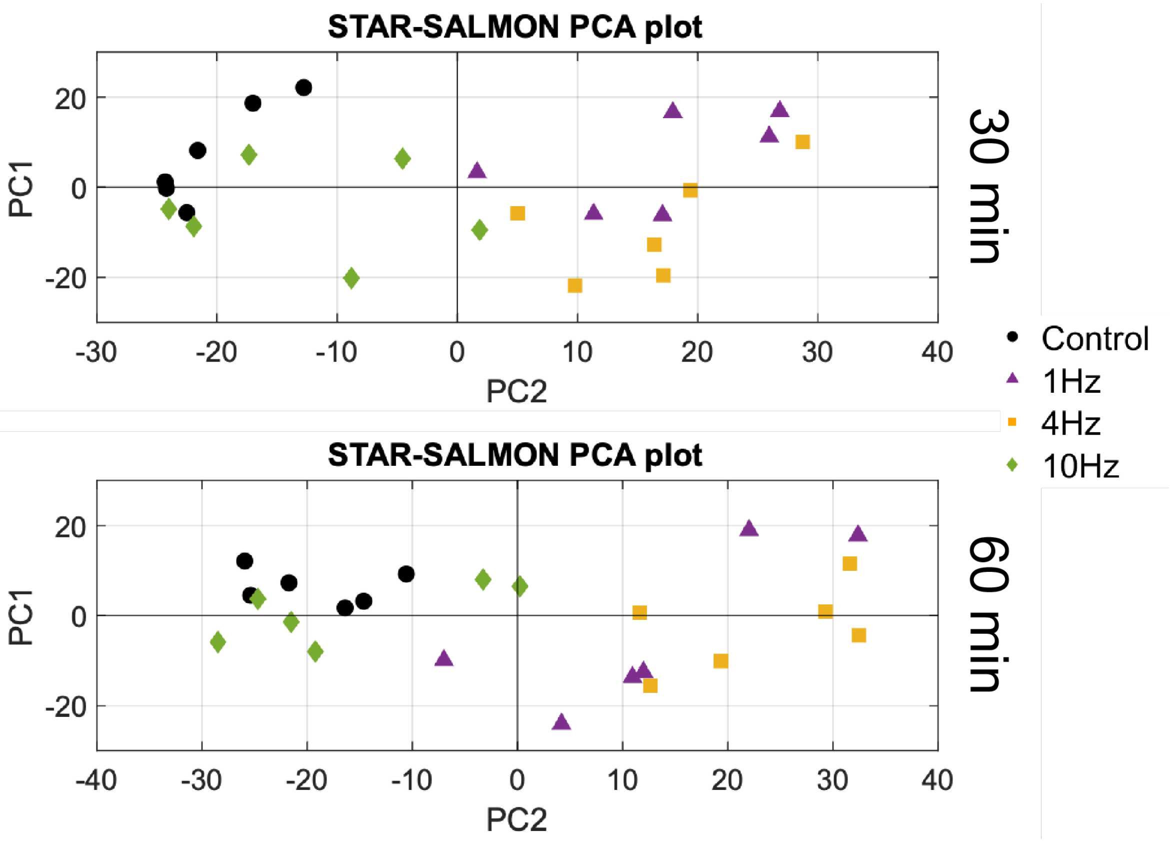


**Figure S2:** Principal component analysis (PCA) plots for 30 min and 60 min exercise experiments.

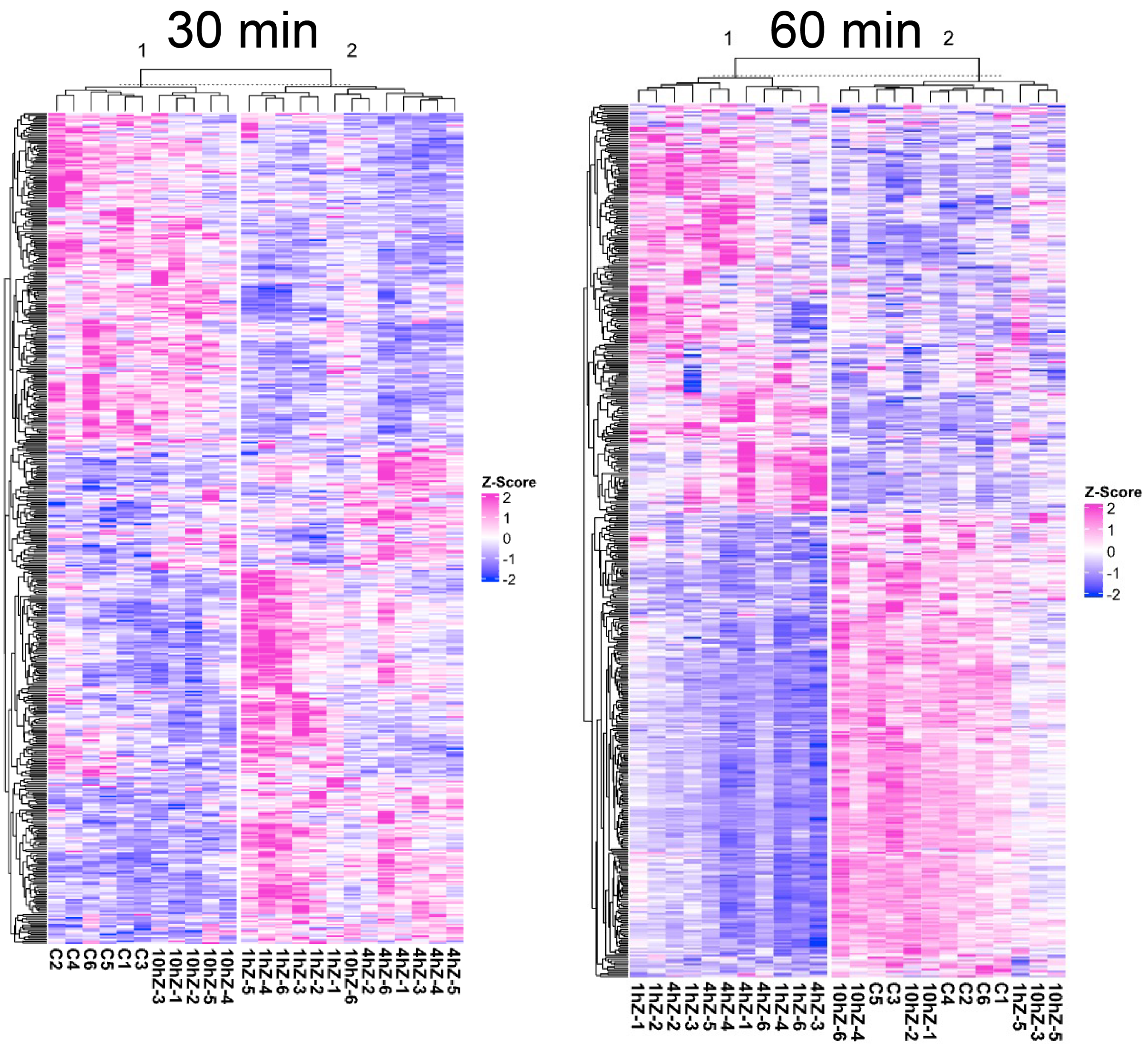


**Figure S3:** RNA-seq heatmaps. Heatmaps show top 500 differentially expressed genes. Genes were using hierarchical clustering on a z-score normalized matrix of transcripts per million.

**Link to Supplemental Movie S1 and S2.**
